# SGLT2 inhibitors protect against diabetic cardiomyopathy and atrial fibrillation through a CaMKII independent mechanism

**DOI:** 10.1101/2024.09.23.614368

**Authors:** Alex Severino, Oscar E Reyes-Gaido, Pauline Nguyen, Ahmed Elkarim, Elizabeth D Luczak, Olurotimi O. Mesubi

## Abstract

Ca^2+^/calmodulin-dependent protein kinase II (CaMKII) has been implicated as an important mediator of the increasingly evident cardioprotective benefits exerted by sodium– glucose transport protein 2 channel inhibitors (SGLT2i). However, the exact nature of the relationship between CaMKII and SGLT2i remains unclear. Here, we find that empagliflozin but not dapagliflozin attenuated susceptibility to atrial fibrillation (AF) in a type 2 diabetic (T2D) mouse model. However, both empagliflozin and dapagliflozin protected from diabetic cardiomyopathy in T2D mice. We then used real-time microscopy of neonatal rat ventricular cardiomyocytes (NRVMs) with the CaMKII biosensor - CaMKAR to demonstrate that direct inhibition of CaMKII is not essential for the effects of SGLT2i in these cells. Therefore, we conclude that the benefits of SGLT2i in heart disease likely occur through indirect modulation of CaMKII activity, or possibly through an alternative pathway altogether.

## Introduction

Sodium-glucose transport protein 2 channel inhibitors (SGLT2i) have emerged as an important class of pharmaceuticals in the management of cardiac disease. Initially intended as agents for glycemic control in patients with type 2 diabetes mellitus (T2D), these inhibitors have subsequently been shown to improve cardiovascular outcomes in patients with heart failure independent of comorbid diabetes (1–3). SGLT2i are also under continued investigation for their potential to protect against arrhythmias. Specifically, broad post-hoc meta-analysis of clinical trials has demonstrated that treatment with empagliflozin or dapagliflozin, among other SGLT2i, significantly reduces the risk of atrial fibrillation (AF) in patients with T2D, heart failure, or chronic kidney disease (4).

Despite the substantive fund of clinical evidence supporting the cardioprotective effects of these drugs, a precise mechanistic understanding of their role in the heart remains unclear. Intriguingly, there is no current evidence that SGLT2 channels are expressed in cardiac tissue (5), and recent work has demonstrated that genetic knockout of the SGLT2 channel in mice is not sufficient to reproduce the amelioration in heart failure with reduced ejection fraction (HFrEF) that is seen with empagliflozin treatment (6). These findings strongly suggest the existence of an unidentified molecular target of SGLT2i in the heart that is mediating these results.

The multifunctional Ca^2+^/calmodulin-dependent protein kinase II (CaMKII) is a key regulator of cardiac physiology and a known driver of heart failure and arrythmias, especially in models of diabetes (7). We have shown that diabetic mice have increased susceptibility toward AF that requires the presence of CaMKII (8). CaMKII has been proposed as a mediator in the action of SGLT2i against AF and cardiac injury (9–12).

Here, we sought to determine whether the cardiovascular therapeutic benefits of SGLT2i - empagliflozin and/or dapagliflozin are due, at least in part, to CaMKII inhibition. We evaluated responses to empagliflozin and dapagliflozin in an established T2D mouse model where excessive CaMKII activation is known to contribute to myocardial dysfunction and enhanced inducibility of AF (8). We used a new high performance CaMKII Activity Reporter (CaMKAR) recently developed in our laboratory (13) to quantify the activity of CaMKII in neonatal rat ventricular cardiomyocytes after administration of either empagliflozin or dapagliflozin.

## Methods

### Animal Care

All mouse experiments were carried out in accordance with the Guide for the Care and Use of Laboratory Animals per the US National Institutes of Health (NIH Publication 8^th^ edition, update 2011) under protocols approved by the Johns Hopkins University Animal Care and Use Committee. Experimental studies were performed on male mice on a C57BL/6J background purchased from The Jackson Laboratory (ME, USA). The mice were housed in a facility with 12-hour light/12-hour dark cycle at 22 ± 1 °C and 40 ± 10% humidity.

### Mouse Model

T2D mice and littermate controls were generated as previously described (8) with the following modifications. In brief, to induce the T2D mouse model, male diet-induced obese (DIO) C57BL/6J mice and controls placed on diet at six weeks of age were purchased from The Jackson Laboratory (Bar Harbor, ME, USA) after five weeks of initiation of diet. The DIO mice were then maintained on a high-fat diet (HFD – 60 kcal% fat – D12492, Research Diets, New Brunswick, NJ, USA) while the control mice were maintained on normal chow diet (NCD – 7913 irradiated NIH-31 modified 6% mouse/rat diet – 15 kg, Envigo, Indianapolis, IN). The DIO mice received daily intraperitoneal (i.p.) injections of low dose streptozotocin (STZ) (40mg/kg/day, Sigma-Aldrich) dissolved in a citrate buffer (citric acid and sodium citrate, pH 4.0) for three consecutive days after a six hour fast on each day. The control mice received daily i.p. injections of citrate buffer for three consecutive days. Blood glucose via tail vein using a glucometer (OneTouch Ultra 2 meter), and body weight were checked one week after STZ or citrate buffer injections. Mice were maintained on the respective diets for 9 – 11 weeks following i.p. injections.

### SGLT2i therapy

The mice were divided into four groups: control group (non-diabetic mice) treated with vehicle control - 0.5% hydroxypropyl methylcellulose (HPMC) in drinking water, T2D, T2D mice treated with 30mg/kg/day empagliflozin dissolved in 0.5% HPMC in drinking water (T2D + Empa) and T2D mice treated with 5mg/kg/day dapagliflozin dissolved in 0.5% HPMC in drinking water (T2D + Dapa). The doses of empagliflozin and dapagliflozin were based on prior studies (14) shown to correspond to effective cardiovascular therapeutic doses used in humans (15). Treatment with vehicle, empagliflozin or dapagliflozin was initiated one week after i.p. injections and maintained for 10 – 11 weeks. Empagliflozin (Cat #HY-15409) and dapagliflozin (Cat #HY-10450) were purchased from MedChemExpress (Monmouth Junction, NJ, USA).

### In-vivo Electrophysiologic and Atrial Fibrillation Studies

Susceptibility to AF was assessed in mice by rapid atrial burst pacing from the right atrium of anesthetized mice during invasive electrophysiologic studies as previously described (8,16–19). A total of five burst pacing rounds was performed in each mouse and a mouse was defined as inducible for AF if 1 or more bursts (out of 5) resulted in AF. We also assessed AF susceptibility using a definition of AF inducibility as 2 or more bursts (out of 5).

### Echocardiography

Cardiac function and dimensions were assessed by transthoracic M-mode echocardiography in conscious mice using the Sequoia C256 ultrasound system (Malvern, PA) equipped with a 15-MHz linear transducer as previously described (20). Measurements were averaged over 3 to 5 beats at physiological heart rates. An experienced blinded operator performed image acquisition and analysis.

### Cell Culture

Neonatal rat ventricular cardiomyocytes (NRVMs) were obtained as previously described (21) from day 1-2 Sprague-Dawley rats and maintained in DMEM (L-glutamine, Sodium Pyruvate, Non-essential amino acids; Gibco) supplemented with 2% FBS (Gibco) and Pen/Strep (Gibco) at a density of 500,000 cells/mL. After isolation, cells were maintained in a humidified incubator at 37C with 5% CO_2_.

### Gene Transfer in Cultured Cells

Lentivirus promoting CaMKAR expression was produced as previously described (13) and stored in aliquots at −80C until ready for use. NRVMs were infected 1 day after isolation with a media change into DMEM with 2% FBS containing lentivirus at the indicated multiplicity of infection for at least 48 hours.

### Microscopy

Timelapse microscopy of CaMKAR fluorescence was performed using materials and techniques previously described (13). In short, imaging was conducted using an Olympus IX-83 inverted widefield microscope along with a Lumencor SOLA light source. CaMKAR fluorescence was captured by an ORCA Flash 4.0 camera using 200 ms exposure through dual excitation filters (ET402/15x and ET 490/20x) as well as a single emission filter (ET 525/35m) supplied by Chroma Technology. CaMKAR signal (R) was defined as the ratio of 488nm-induced to 405nm-induced fluorescence. At the time of imaging, culture medium was removed from cells and replaced with imaging buffer (Tyrode’s salts solution Sigma-Aldrich Cat #T2397, 10mM glucose, 1.8mM CaCl). In order to ensure electrical capture during pacing, calcium signal was simultaneously measured using Calbryte 590 AM dye (AAT Bioquest Cat #20700) with fluorescence viewed through an mCherry 587/610 excitation and emission filter. Calbryte dye was resuspended in DMSO before diluting in imaging buffer and subsequently added to NRVMs for a final concentration of 5 μM. Dye was incubated in a 37 C, 5% CO_2_ incubator for 30 minutes before pacing. In order to investigate the effects of SGLT2i in this system, cells were preincubated with either empagliflozin or dapagliflozin (MedChemExpress Cat # HY-15409 and Cat # HY-10450, Adooq Bioscience Cat #A12440 and Cat #A10285) for the specified amount of time before imaging. AS100397 (10 μM) and Calyculin A (Cayman Chemical Company Cat #19246) (100nm) were added to specified wells 10 minutes before imaging. Using computational methods previously described (13),Otsu segmentation was used to track individual cells and their fluorescence intensity in 488 nm and 405 nm channels at each timepoint (every 5 seconds) in CellProfiler, and image analysis was conducted using custom scripts in Fiji/ImageJ. Statistical tests and graphical representation were performed in GraphPad Prism 10.

### Calcium Imaging and Analysis

Ca^2+^ transients were measured using a previously described custom IonOptix system (22). NRVMs were loaded with 2 μM fura-2-AM (Invitrogen Cat #F1221) reconstituted in DMSO and dissolved in Tyrode’s salts solution, and cells were then incubated at room temperature for 20 minutes. Cells underwent excitation at 340 and 380 nm at 250 Hz with 510 nm wavelength emission measured using a single photon multiplier (PMT). Cells were paced at twice minimum capture threshold while recordings were collected from single cells over the course of 15 to 25 transients. Transient data from each cell were modeled using the CytoSolver 3.0 Transient Analysis software and averaged for comparison across treatment conditions. Cells were preincubated with empagliflozin or dapagliflozin for the specified amount of time, or with AS100397 inhibitor for 10 minutes before transient measurements. Statistical tests and graphical representation were performed in GraphPad Prism 10.

## Results

### T2D mouse model responses to empagliflozin and dapagliflozin

In order to investigate the potential role of CaMKII in the cardioprotective effects of SGLT2 inhibition we deployed a model of T2D where CaMKII is established to promote myocardial dysfunction and AF induction. We induced T2D in mice by feeding them a high-fat diet (60% kcal) and administering low-dose streptozotocin (STZ) (see Methods) (23). We then initiated treatment with either empagliflozin, dapagliflozin, or vehicle in the drinking water for 10-11 weeks and measured the inducibility of AF, along with measurements of blood glucose and echocardiography (Fig 1a). T2D mice demonstrated increased blood glucose levels and higher body weights compared to non-diabetic littermate controls 1 week after STZ administration and at the time of cardiac studies (Fig 1b).

**Figure 1.**
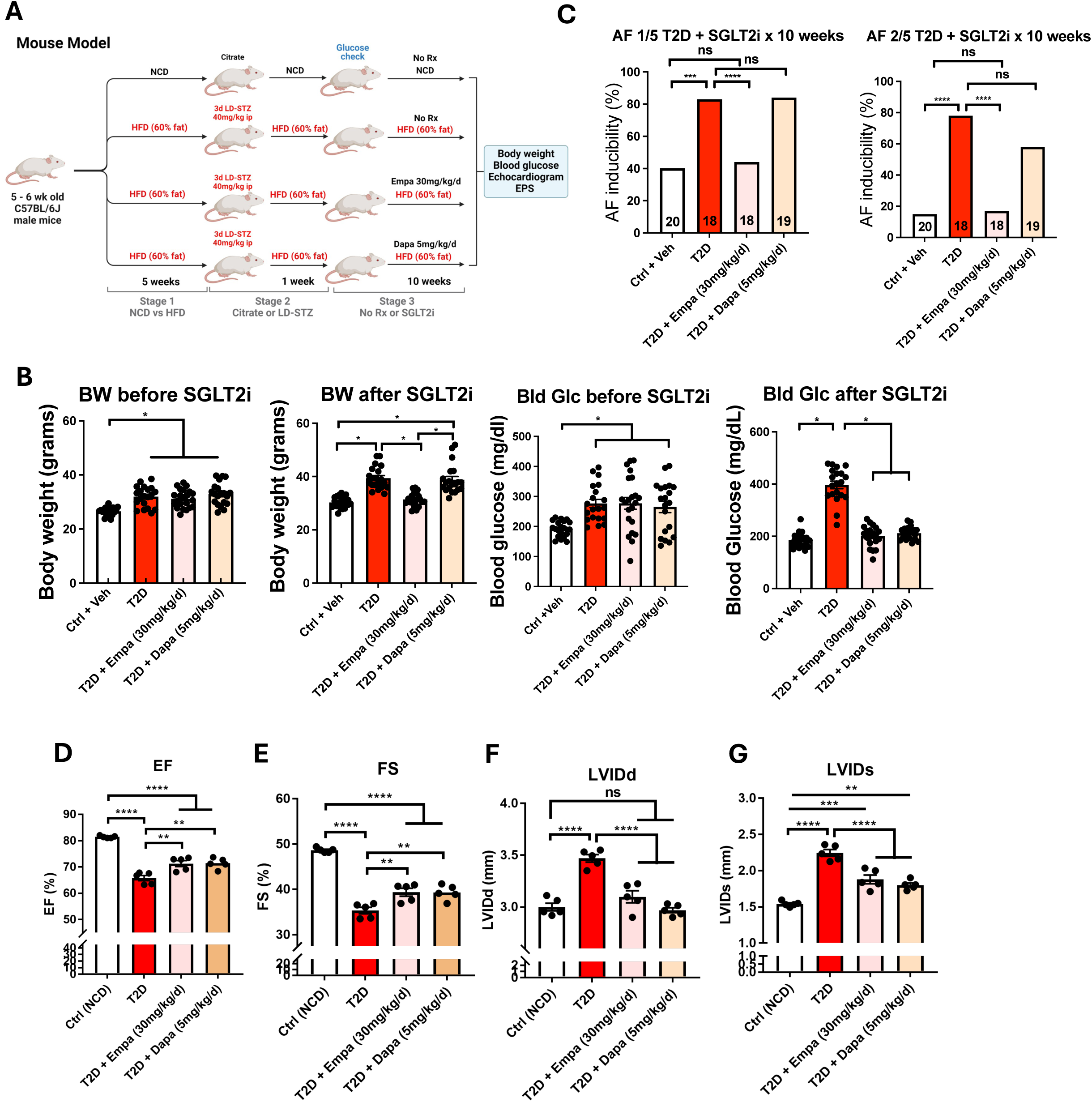
Effects of empagliflozin and dapagliflozin on diabetic cardiomyopathy in mice. (A) Mice were given either normal chow (NCD) or high-fat diet (HFD) and exposed to either streptozotocin (STZ) or vehicle (citrate). Mice were subsequently given empagliflozin, dapagliflozin, or vehicle water before echocardiogram and electrophysiologic studies (EPS). (B) Body weight (BW) and blood glucose (Bld Glc) measurements of mice before intervention (left) and at time of EPS. (C) Inducibility of AF in T2D mice after 10-week treatment with either empagliflozin or dapagliflozin where AF is defined as 1/5 induced episodes (left) or 2/5 induced episodes (right). (D-G) Echocardiographic parameters of cardiomyopathy in T2D mice treated with either empagliflozin or dapagliflozin for 10 weeks. ns, *P < 0.05, **P < 0.005, ***P < 0.0005, ****P < 0.0001. Significance was determined by χ^2^ test (C) and ordinary one-way ANOVA with multiple comparisons (B, D-G).

The T2D mice had a significant increase in the inducibility of AF compared to non-diabetic controls (40% controls vs 83% T2D by AF 1/5, *p* = 0.0002), a phenotype that was rescued with treatment by empagliflozin (44% empagliflozin vs 40% controls by AF 1/5, p = 0.7003) but not dapagliflozin (84% dapagliflozin vs 40 % controls by AF 1/5, p < 0.0001) (Fig 1c). These findings remained consistent whether AF inducibility was defined as 1/5 or 2/5 episodes of AF induction (see Methods) (Fig 1c). Both the empagliflozin and dapagliflozin treatment groups exhibited normal blood glucose levels similar to non-diabetic controls at the conclusion of the study, consistent with the ability of SGLT2i to restore glycemic control in the setting of diabetes and supporting the notion of successful drug delivery (Fig 1b). Of note, the body weights of empagliflozin-treated mice were equivalent to those of control mice at the end of 10 weeks, while the mice receiving dapagliflozin exhibited similar weight gain to the untreated T2D group (Fig 1b). Both treatment groups demonstrated improved echocardiographic parameters such as ejection fraction, fractional shortening, and left ventricular wall thickness compared to untreated T2D mice (Fig 1d-g).

### Lack of CaMKII inhibition in empagliflozin or dapagliflozin treated cardiomyocytes

Our in vivo studies showed consistent metabolic and myocardial performance benefits for empagliflozin and dapagliflozin treatment in T2D. In contrast, the responses to empagliflozin and dapaglifozin in AF inducibility for mice with T2D were inconsistent. Collectively, we interpreted these findings as being inconclusive for supporting a role of myocardial CaMKII inhibition as part of the mechanism for empagliflozin and dapagliflozin. In order to directly address this knowledge gap we utilized CaMKAR, a recently developed enhanced green fluorescent protein (eGFP)-based biosensor for CaMKII activity that is deployable in neonatal rat ventricular myocytes (NRVMs) (24). NRVMs made to express CaMKAR through adenoviral infection (see Methods) were incubated with either empagliflozin or dapagliflozin and then electrically paced under a fluorescence microscope to assess real-time levels of CaMKII activation. The area under the curve (AUC) of the CaMKAR fluorescence tracings was used as a readout for CaMKII activity in order to evaluate for enzymatic inhibition compared to controls (Fig 2a), as described (see Methods) (13).

**Figure 2.**
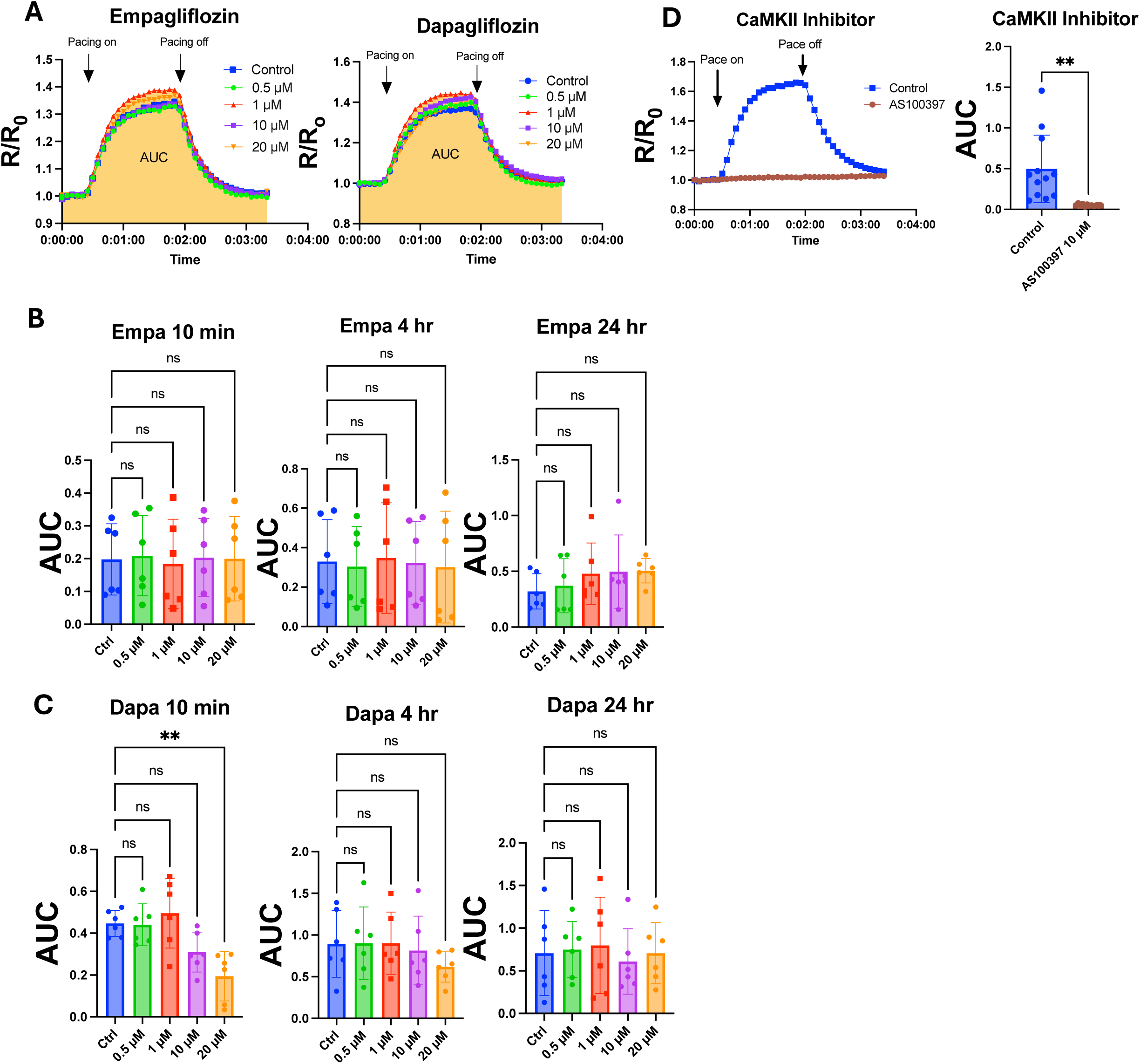
CaMKII activity in SGLT2i-treated neonatal rat ventricular cardiomyocytes (NRVMs) measured by CaMKAR. (A) Representative images of time-lapse fluorescence tracings using CaMKAR in SGLT2i-treated NRVMs during electrical stimulation at 2 Hz. Quantification of area-under-the-curve (AUC) for time-lapse recordings of NRVMs after acute and chronic treatment with empagliflozin (B) or dapagliflozin (C) compared to untreated cells. (D) Representative tracing and AUC quantification for NRVMs treated with the known CaMKII inhibitor AS100397 (10 µM) compared to controls during electrical stimulation. All observations are aggregated from fluorescence of many cells in field of view. ns, **P < 0.005. Significance was determined by ordinary one-way ANOVA with multiple comparisons (B,C) and Student’s t-test (D).

We incubated NRVMs with SGLT2i for either 10 minutes, 4 hours, or 24 hours before imaging to determine whether there was a difference between acute and chronic drug treatment. Across all timepoints, we observed no significant difference in CaMKAR fluorescence when cells were treated with 1 μM empagliflozin or dapagliflozin, a concentration equivalent to maximum plasma drug levels achieved in vivo (15) (Fig 2b, 2c). Alternatively, cells pretreated with the known CaMKII inhibitor AS100397 exhibited significantly decreased CaMKAR signal in this system (p = 0.003) (Fig 2d). For empagliflozin, no significant inhibition of CaMKII was detected up to a treatment concentration of 20 μM (Fig 2b). Interestingly, there was an observed decrease in CaMKII activity with 20 μM dapagliflozin treatment for 10 minutes (*p* = 0.006), although this effect disappeared when cells were incubated for 4 hours or 24 hours before imaging (Fig 2c). Taken together these data do not support the view that empagliflozin or dapagliflozin inhibit myocardial CaMKII at therapeutically relevant concentrations.

### Empagliflozin and dapagliflozin actions on intracellular Ca^2+^ transients

As a Ca^2+^ sensor, activation of CaMKII is tightly coupled to the morphology and frequency of calcium transients during cardiac depolarization (25). Furthermore, CaMKII activity affects multiple proteins important for regulating intracellular Ca^2+^ concentration. Based on these considerations we next measured the effects of empagliflozin and dapagliflozin on intracellular Ca^2+^ transients in NRVMs using fura-2 AM (see Methods). With both empagliflozin and dapagliflozin, the amplitude of Ca^2+^ transients was increased after 10 minutes of treatment at 1 μM (0.38 fu in DMSO vs 0.57 fu in empagliflozin (*p* < 0.0001) vs 0.50 fu in dapagliflozin (p < 0.0001)) (Fig 3a). In contrast, 10-minute treatment of NRVMs with AS100397 led to a significantly decreased amplitude of measured Ca^2+^ transients compared to DMSO controls, as anticipated in response to CaMKII inhibition (Fig 3b). We also measured the time constant, tau, of calcium transients (Fig 3c) and found that 4-hour treatment with empagliflozin and dapagliflozin significantly increased time constant compared to controls (0.22 seconds control vs 0.31 seconds empagliflozin vs 0.34 seconds dapagliflozin, p = 0.0125, 0.0005). The AS100397 condition showed no significant change in transient time constant compared to control cells (Fig 3b). Time to half-peak (t_1/2_-upstroke) was elevated with AS100397 but showed discordant effects in both empagliflozin and dapagliflozin treatments without a clear dose-response pattern across treatment durations (Fig 3d, 3e).

**Figure 3.**
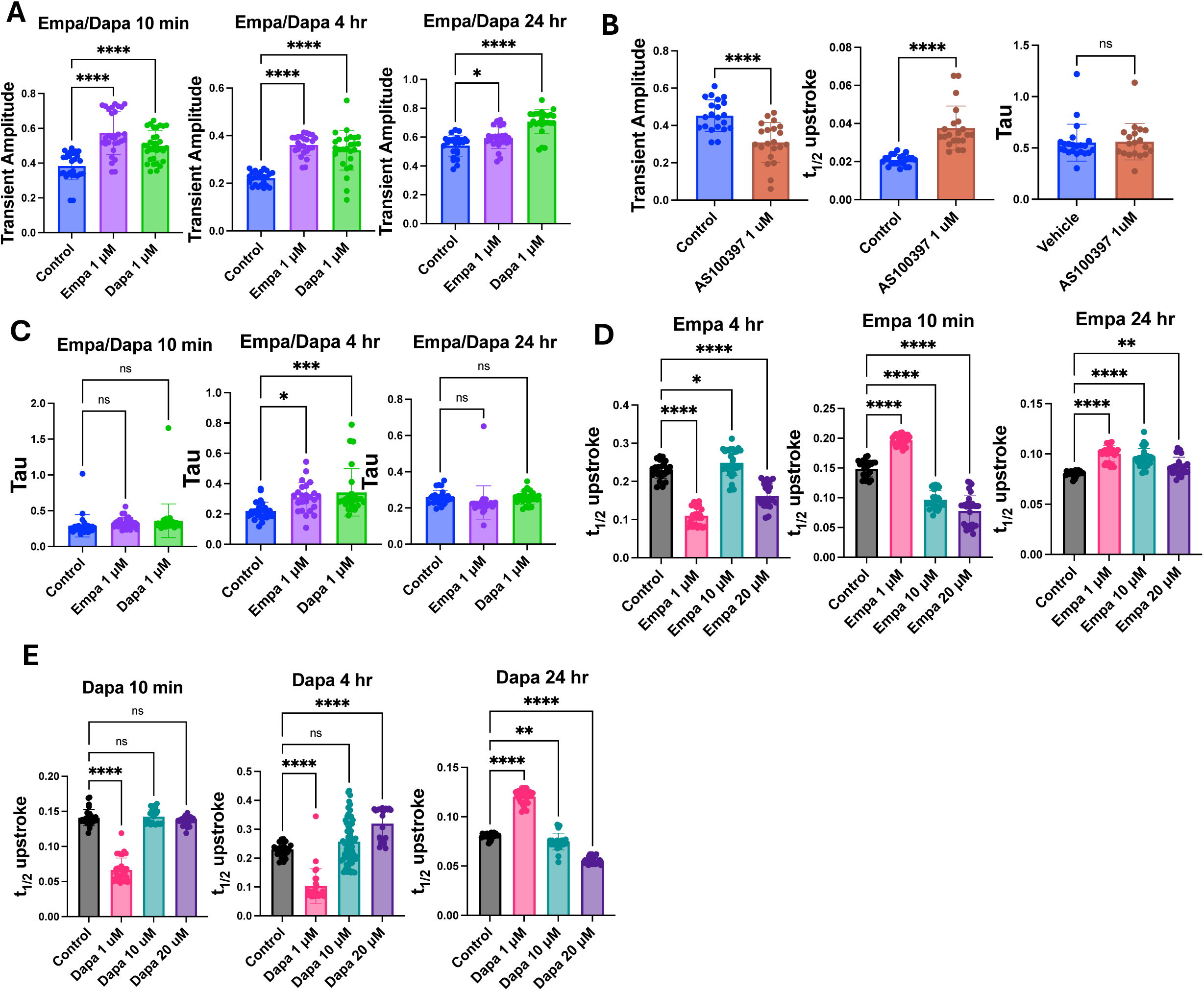
Empagliflozin and dapagliflozin alter calcium handling in electrically-paced NRVMs. (A) Amplitude of calcium transients in NRVMs paced at 2 Hz. (B) Amplitude of calcium transients, time constant (tau), and time to half-peak (t_1/2_ upstroke) in paced NRVMs pretreated for 10 minutes with the CaMKII inhibitor AS100397 (1 µM) compared to controls. (C) Tau measurements for empagliflozin and dapagliflozin treatments compared to controls after acute and chronic exposure. Time to half-peak for empagliflozin (D) and dapagliflozin (E) after increasing treatment duration and concentration. All observations are readings from single cells, each measured for at least 10 seconds. *P < 0.05, **P < 0.005, ***P < 0.0005, ****P < 0.0001. Significance was determined by ordinary one-way ANOVA with multiple comparisons (A, C-E) or Student’s t-test (B).

## Discussion

CaMKII inhibition has emerged as an important candidate action for the positive effects of SGLT2i in cardiac disease. Previous studies have demonstrated an inhibitory effect of SGLT2i on cardiac sodium channels, resulting in attenuation of sodium influx through the Na^+^/H^+^ exchanger in cardiomyocytes (26). It has been suggested that this reduction in intracellular sodium alters the equilibrium of sodium-calcium exchanger (NCX) channels, thereby reducing intracellular calcium levels and dampening CaMKII activation. CaMKII has also been shown to be necessary for inhibition of the late sodium current in human cardiomyocytes (27). However, direct evidence confirming inhibition of CaMKII as the linchpin of SGLT2i-induced benefits in the heart is lacking, and a validated molecular target for these drugs in cardiac tissue remains unclear. Indirect markers of decreased activation have been used to support the CaMKII hypothesis, such as altered CaMKII pulldown with downstream targets (28) and reduced phosphorylation of the ryanodine receptor (29).

Here, our results using CaMKAR as a direct readout of CaMKII activation does not indicate that treatment of NRVMs with empagliflozin nor dapagliflozin generates a significant reduction in CaMKII activity upon electrical pacing. Other work has demonstrated a time-dependence of empagliflozin’s effects in cardiomyocytes (28), prompting us to examine a wide range of timepoints for SGLT2i treatment. We observed no significant difference in CaMKAR activation compared to controls after both acute (10 minute) and chronic (4 and 24 hour) treatment. We do note a statistically significant inhibitory effect on CaMKII in the acute treatment condition with dapagliflozin at 20μM. Because this inhibition was only observed at a concentration much greater than the accepted physiologic drug levels achieved in plasma and did not persist with longer exposure, we are hesitant to conclude that this finding is representative of a phenomenon occurring in patients taking these medications.

We do observe that SGLT2i treatment using concentrations that approximate physiologic dosing increases Ca^2+^ transient amplitude, a sign of altered calcium handling and positive inotropy (30). Discordantly, treatment with the known CaMKII inhibitor AS100397 gives a predicted response of diminishing the calcium transient. This is exhibited by decreased transient amplitudes and an increase in the t_1/2_-peak, although we did not observe a significant change in the time constant from AS100397 treatment. Of note, empagliflozin and dapagliflozin did increase the time constant after 4 hours but not after 10 minutes or 24 hours which is suggestive of slowed calcium waves and also highlights how time-dependent the actions of SGLT2i appear to be. Effects on t_1/2_-peak showed variation with different concentrations and treatment durations of SGLT2i without creating a clear signal. Collectively, the distinct differences in the profiles of Ca^2+^ transients between SGLT2i and AS100397 conditions further indicate that empagliflozin and dapagliflozin appear to be influencing intracellular calcium handling in a manner distinct from direct CaMKII inhibition.

We acknowledge that the models used in our experiments are not without limitations. For instance, most clinical trials that have demonstrated features of cardioprotection from SGLT2i consisted of patients from the heart failure population (2,31–33). The concept of diseased myocardium is not captured in our use of NRVMs, which are healthy neonatal cells, while the T2D mice had evidence of some cardiomyopathy and systolic dysfunction. Other published experiments have utilized models that attempted to incorporate the presence of pathology in cardiomyocytes prior to administration of SGLT2i, such as by isolating adult cardiomyocytes from mice undergoing transverse aortic constriction (TAC) (34,35). It is therefore conceivable that cardiomyocytes challenged with ischemia or other such metabolic stressors undergo alterations in cellular homeostatic and ion-handling mechanisms compared to healthy neonatal cells that are necessary for SGLT2i to inhibit CaMKII activity. However, we have shown here that empagliflozin and dapagliflozin do exert changes in intracellular calcium signaling during pacing of NRVMs. This finding reassures us that SGTL2i are not inert in our model and therefore could conceivably be expected to show a signal of CaMKII inhibition if indeed one existed.

SGLT2i are becoming an increasingly employed cornerstone of therapeutic regimens for cardiac disease due to post-hoc evidence from clinical trials demonstrating cardioprotective effects. Our results contribute new evidence that empagliflozin protects against atrial fibrillation inducibility in mice, which encouragingly recapitulates in our model the antiarrhythmic signal observed in clinical trials. The dissonance between atrial fibrillation protection by empagliflozin but not dapagliflozin suggests the possibility that the observed effects of these drugs in patients might not be attributable to a class effect but rather to distinct mechanisms. In congruence with these results, a recent multicenter retrospective cohort study in patients with heart failure found that empagliflozin may be superior to dapagliflozin in preventing all-cause mortality and hospitalization (36).

To our knowledge, our results here are the first to use such a direct measurement of CaMKII activity afforded by the biosensor CaMKAR in order to assess for CaMKII inhibition by SGLT2i. While we concur with previous studies that SGLT2i are active compounds in cardiomyocytes which are capable of promoting changes in intracellular ion homeostasis independently of SGLT2 channels, we do not find evidence that a decrease in CaMKII activity is the ultimate effector in the pathway by which SGLT2i might be cardioprotective. As the clinical evidence continues to grow suggesting that SGLT2i can improve outcomes in heart failure and arrhythmias, it becomes more crucial than ever to advance our understanding of how these therapeutics are working mechanistically in these circumstances. Identification of a precise molecular target for SGLT2i in the heart will prove critical not only in predicting which patient populations are likely to receive the greatest benefit from their usage in the pursuit of increasingly personalized medicine, but it may also help to guide the path for development of future potent treatment strategies in the management of cardiovascular disease.

## Abbreviations

AF: Atrial fibrillation
CaMKII: Calcium and calmodulin-dependent protein kinase II
CAMKAR: CaMKII Activity Reporter
DIO: Diet-induced obese
DM: Diabetes mellitus
HFD: High fat diet
HPMCH: ydroxypropyl methylcellulose
NRVMs: Neonatal rat ventricular cardiomyocytes
NCD: Normal chow diet
STZ: Streptozocin
T2D: Type 2 diabetes mellitus

## Acknowledgements

We are thankful to the Johns Hopkins Dean’s Year of Research Program for supporting AS, Fondation Leducq for providing the AS100397 compound, Mark Anderson for discussions and review of the manuscript, D. A. Kass and F. Wu for providing NRVMs, Nadan Wang for performing the mouse echocardiograms and Jinying Yang for her assistance in maintaining mouse colonies.

